# Improved freshwater macroinvertebrate detection from eDNA through minimized non-target amplification

**DOI:** 10.1101/2020.04.27.063545

**Authors:** Florian Leese, Mandy Sander, Dominik Buchner, Vasco Elbrecht, Peter Haase, Vera M.A. Zizka

## Abstract

DNA metabarcoding of freshwater communities typically relies on PCR amplification of a fragment of the mitochondrial cytochrome c oxidase (COI) gene with degenerate primers. The advantage of COI is its taxonomic resolution and the availability of an extensive reference database. However, when universal primers are used on environmental DNA (eDNA) isolated from stream water, macroinvertebrate read and OTU numbers are typically “watered down”, i.e. diluted, compared to whole specimen ‘bulk samples’ due to greater co-amplification of abundant non-target taxa such as algae and bacteria. Because stream macroinvertebrate taxa are of prime importance for regulatory biomonitoring, more effective ways to capture their diversity via eDNA isolated from water are important. In this study, we aimed to improve macroinvertebrate assessment from eDNA by minimizing non-target amplification. Therefore, we generated data using universal primers BF2/BR2 throughout 15 months from a German Long-Term Ecological Research (LTER) site, the River Kinzig, to identify most abundant non-target taxa. Based on these data, we designed a new reverse primer (EPTDr2n) with 3’-specificity towards macrozoobenthic taxa and validated its specificity *in silico* together with universal forward primer fwhF2 using available data from GenBank and BOLD. We then performed *in vitro* tests using 20 eDNA samples taken in the Kinzig catchment. We found that the percentage of target reads was much higher for the new primer combination compared to two universal macrozoobenthic primer pairs, BF2/BR2 and fwhF2/fwhR2n (>99 % vs. 21.4 % and 41.25 %, respectively). Likewise, number of detected macroinvertebrate taxa was substantially higher (351 vs. 46 and 170, respectively) and exceeded the number of 257 taxa identified by expert taxonomists at nearby sites across two decades of sampling. While few taxa such as Turbellaria were not detected, we show that the optimized primer avoids the dilution problem and thus significantly improves macroinvertebrate detection for bioassessment and -monitoring.

## Introduction

With environmental DNA (eDNA) extracted from water, aquatic biodiversity can be detected across the tree of life with minimal effort (Taberlet et al. 2018, Grey et al. 2018, Li et al. 2018, Bagley et al. 2019, Zhang et al. 2020). The potential of eDNA methods, in particular eDNA metabarcoding, for aquatic biodiversity and ecological quality assessments is therefore undisputed (Keck et al. 2017, Pawlowski et al. 2018). Especially for amphibians and fish, environmental DNA has been reported to capture species diversity in much greater detail than traditional techniques (Thomsen et al. 2012, Valentini et al. 2016, Stoeckle et al. 2017, Pont et al. 2018, Li et al. 2019). In addition, derived ecological status classes or quality rations (EQRs) based on fish data are consistent between eDNA assessments and traditional techniques (e.g. Hänfling et al. 2016, Pont et al. 2018). Therefore, a roadmap towards the inclusion of eDNA-based methods into regulatory assessment programs is discussed (Pont et al. online early, Leese et al. 2018, Hering et al. 2018, Li et al. 2019). The big advantage of vertebrate biodiversity assessment from eDNA isolated from water, is that primers targeting vertebrate-specific, conserved regions of the mitochondrial 12S or 16S rDNA are typically used providing the majority of reads for the target taxa under study.

While eDNA-based assessments for fish and amphibians are in principle at the level of routine application, the situation for macroinvertebrate species assessments lags behind substantially (Belle et al. 2019, Blackman et al. 2019). One main reason for this is that, in contrast to fish and amphibian eDNA analyses, the typically used metabarcoding fragment is the mitochondrial cytochrome c oxidase I (COI) gene. The big advantage of COI is its taxonomic resolution and the availability of extensive reference data (Ratnasingham and Hebert 2007, Clarke et al. 2017, Andújar et al. 2018, Weigand et al. 2019). However, in contrast to 12S or 16S, COI as a protein-coding gene shows codon degeneracy that limits taxon-specific primer design (Clarke et al. 2014, Deagle et al. 2014, Sharma and Kobayashi 2014) making it almost impossible so far to design really target-specific primers for diverse groups such as ‘invertebrates’ but also fish and amphibians. Elbrecht and Leese (2017a) proposed an approach to consider taxa of a region or only specific taxon groups degenerate primers, depending on the research or applied question. It should be noted, however, that primer degeneracy is no main concern when metabarcoding is used for bulk tissue samples, as this ‘biodiversity soup’ (Yu et al. 2012) mostly contains DNA of the target organisms (Elbrecht et al. 2016, 2017). It becomes a big challenge when trying to detect trace amounts of target DNA in pools of non-target DNA and this is the case with eDNA extracted from water. Samples are dominated by DNA from living bacteria, fungi, and other non-target DNA. When the vast majority of DNA in the sample is non-target DNA, universal primers for COI will amplify a majority of COI fragments from non-target organisms even if primers do not match well. This leads to DNA of the target macroinvertebrates becoming ‘watered down’ (Hajibabaei et al. 2019a). This phenomenon has been reported throughout studies using COI for macroinvertebrate assessments from eDNA. For example, Deiner et al. (2016) reported a large diversity of macroinvertebrate families detected from stream eDNA, yet, small non-target taxa such as Rotifera captured the vast majority of reads. In a study comparing bulk sample and eDNA metabarcoding in New Zealand streams, Macher et al. (2018) detected a vast amount of taxa using eDNA, yet only 21% of reads were assigned to Metazoa. When considering species (>97 % identity) for the important bioindicator taxa Ephemeroptera, Plecoptera, Trichoptera, and Diptera (EPTD), number of reads only comprised 0.6 % compared to about 30 % in bulk samples (Macher et al. 2018, Macher pers. comm.). Similarly, Beentjes et al. (2019) performed eDNA metabarcoding on water from ponds and recovered a majority of non-target taxa reads and OTUs (35.7 % of reads Metazoa, 14% of the OTUs). As in Deiner et al. (2016) and Macher et al. (2018), most of these were meiofaunal metazoans and only a minor faction (mostly < 5 %) were macroinvertebrate insects (see Beentjes et al. 2019 Supp. Fig. S3). Thus, especially for indicator taxa, species or OTUs diversity was low. Li et al. (2018) In the study by Hajibabaei et al. (2019), which coined the term ‘watered-down biodiversity’, number of reads assigned to benthic invertebrates per eDNA sample was two orders of magnitude lower and diversity, measured as the number of exact sequence variants (ESVs) per sample, one order of magnitude lower compared to bulk samples. Pereira-da-Conceicoa et al. (2019) showed that water samples from South African streams contained less than 10 % reads assigned to targeted macroinvertebrate taxa. As a consequence, while aquatic vertebrate assessments already target eDNA as source material, macroinvertebrate biodiversity assessments still focus on whole organismal samples, i.e. bulk samples, in order to maximize species detection. Several studies demonstrated that DNA metabarcoding data obtained from bulk samples yield robust, powerful, and highly resolved data that can be intercalibrated with current environmental assessment procedures (Aylagas et al. 2014, 2018, Elbrecht et al. 2017, Kuntke et al. 2020). However, collecting and sorting individual macroinvertebrates, is a time-consuming (and thus costly) step that greatly hinders the adoption of the methods at a broader scale (see Blackman et al. 2019 for a discussion). As alternative methods that omit the time-consuming step of specimen-picking, three approaches are proposed: i) using only the preservative liquid (Hajibabaei et al. 2012, Zizka et al. 2018, Martins et al. 2019, Erdozain et al. 2019), ii) using completely homogenised environmental bulk samples without sorting of MZB specimens (Pereira-da-Conceicoa et al. 2019), or iii) using simply a water sample as done for fish or amphibian species (Macher et al. 2018, see Blackman et al. 2019 for a review). As eDNA isolated directly from water is by far the simplest and most economic approach, it is regarded as the ideal solution when the aim is to maximize data generation for aquatic biomonitoring 2.0 (Baird and Hajibabaei 2012, Bush et al. 2017). Therefore, the aim of this study was minimize the effect of target taxon read and diversity dilution in eDNA samples by designing and testing (*in silico* and *in vitro)* a new specific COI eDNA primer targeting macroinvertebrates. We compared the efficiency by comparing proportion of recovered macroinvertebrate reads from eDNA samples as well as number of OTUs for three different primer pairs. To this end, we collected eDNA at multiple stations of a German Long-Term Ecological Research (LTER)(Mirtl et al. 2018) site, the river Kinzig catchment. For this LTER site biodiversity data, especially benthic macroinvertebrate occurrences, have been compiled for over two decades allowing the validation of eDNA results.

## Materials and Methods

### Material

Water samples were obtained from one site at the river Kinzig (Hesse, Germany) within the Long-Term Ecological Research (LTER) site Rhine-Main-Observatory (RMO) from May 2017 to August 2018. LTER sites, as the RMO, are particularly suited for comparative analyses as they provide a wealth of long-term biodiversity data (Haase et al. 2016, Kuemmerlen et al. 2016). In total we collected 102 samples from the Kinzig (site 54, see Fig. 1) in biweekly intervals. Three 1 L-samples were taken per sampling day: surface water in the middle of the stream, 10 cm above riverbed in the middle of the stream, and riverbank. The performance of the newly designed primer was evaluated using a subset of ten of these samples as well as using ten water samples collected further upstream (river Kinzig network). Those additional river Kinzig network water samples were collected from the Kinzig and its tributaries within the RMO in spring 2019 (see Supporting Information S1). After sampling 3x 1 L of stream water for the biweekly monitoring (2017, 2018) and 1x 1 L for the river Kinzig network (2019), samples were stored at −20 °C until further processing. Samples were filtered using a vacuum pump and DNA was captured on 0.45 μm Cellulose Nitrate membrane-filters (diameter 47 mm, Nalgene). Blank/negative controls were included for each filtering day and person that filtered. For filtering, a tube attached to an electric pump was used and for working as sterile as possible, gloves were changed, and the workspace was cleaned with ethanol and bleach (4-5 % sodium hypochlorite) in between samples. Depending on turbidity, one, two, or sometimes three filters were used per sample. Filters were stored in Eppendorf tubes filled with 96 % ethanol. Downstream eDNA processing was carried out in a dedicated eDNA laboratory. Full-body protective equipment was worn, and surfaces were sterilized with UV-light after each work cycle. For DNA extraction, filters were placed in sterile Petri dishes and dried overnight at room temperature. DNA extraction was performed following a salt-precipitation protocol (Weiss and Leese 2016). Next, 1.5 μl RNase A (10 mg/ml) was added to each sample and incubated for 30 minutes at 37 °C and 300 rpm in an Eppendorf ThermoMixer C (Eppendorf AG, Hamburg, Germany). Samples were then cleaned up using a Qiagen MinElute DNeasy Blood & Tissue Kit (Hilden, Germany). Samples were eluted in 30 μl PCR H_2_O.

**Figure 1:**
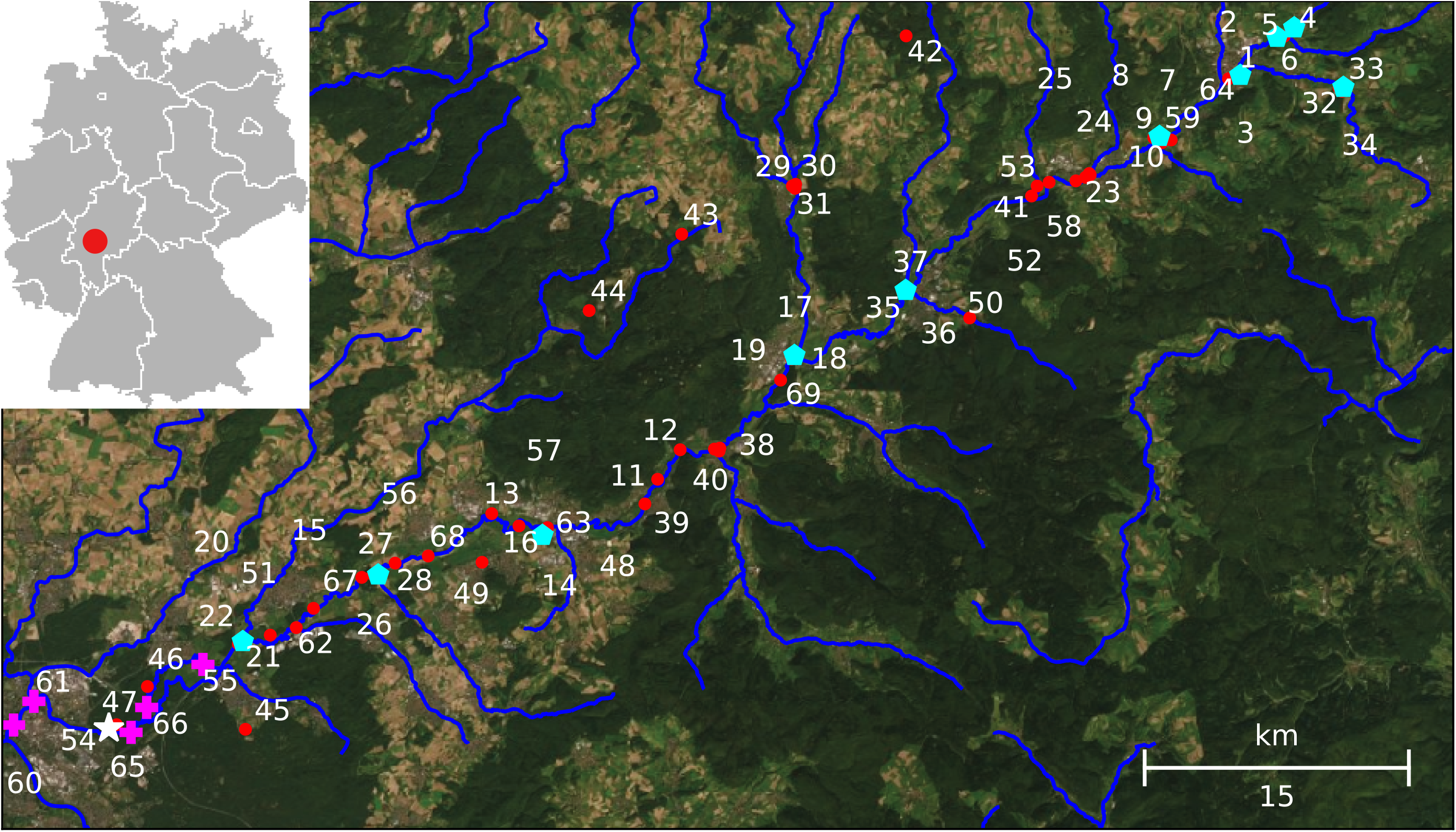
Sampling locations within the Long-Term Ecological Research (LTER) site Rhine-Main-Observatory in Hesse, Germany. The white star (site 54) indicates the biweekly sampling site. Purple crosses indicate long-term (morphological) monitoring sites in close vicinity to site 54 that were used for morphological comparison. Blue pentagons indicate the River Kinzig network samples. Red dots represent other (morphological) monitoring sites. In case of several nearby sampling sites, more than one number is given. See Supplementary Tab. 1 for further details.

### DNA metabarcoding with BF2/BR2 for biweekly data set

We applied a two-step PCR approach for the long-term biweekly data set (Zizka et al. 2019). For the first PCR step, universal BF2/BR2 primers (Elbrecht and Leese 2017a) were used. For the second step, fusion primers with an unique inline shift and Illumina adapters attached to the 5’-end were used. For the first step four PCR replicates were processed per extract and for the second step, two of them were pooled together, resulting in two PCR replicates for each sample. All PCR reactions contained 1x Multiplex PCR Master Mix (Qiagen Multiplex PCR Plus Kit, Qiagen, Hilden, Germany), 0.2 μM of each primer, 0.5 × Q-solution, 1 μl DNA extract/PCR product (concentration not measured) filled up with RNase-free water to a total volume of 50 μl. PCRs were conducted as follows: First step PCRs consisted of 5 min initial denaturation at 95 °C, 30 cycles of 30 s at 95 °C, 90 s at 50 °C and 2 min at 72 °C, followed by a final elongation for 10 min at 68 °C. Second step PCR was identical to first-step PCR only 15 cycles were run with an annealing temperature of 57 °C. The DNA concentration of each PCR product of the second step PCR was quantified on a Fragment Analyzer™ Automated CE System (Advanced Analytical Technologies GmbH, Heidelberg, Germany) using the NGS Standard Sensitivity Kit. PCR products were equimolarly pooled. The pool was purified with a Qiagen MinElute Reaction Cleanup Kit (Hilden, Germany) to remove BSA prior to a 0.76x SPRIselect left-sided selection for BF2/BR2 (Beckman Coulter, CA, USA). Libraries were sequenced on an Illumina MiSeq system with a 250 bp paired-end read kit v2 with 5 % Phi-X spike-in. Sequencing was carried out by CeGaT GmbH (Tübingen, Germany). 102 samples were sequenced for the BF2/BR2 primer pairs from site 54 (biweekly sampling). We obtained between 27,934 and 282,981 reads per sampling site/time point (bioproject no: tbc, Sander et al. unpublished data).

For bioinformatic analysis sequences were assigned to their original sample using JAMP v0.67 (https://github.com/VascoElbrecht/JAMP)(Elbrecht et al. 2018). Standard settings were used to perform subsequent data processing. Paired-end reads were first merged (module U_merge_PE (fastq_maxiffs = 99, for the maximum number of mismatches in the alignment and fastq_pctid = 90, for the minimum %id of the alignment)) and if needed, the reverse complements of the sequences were built (U_revcomp) with usearch v11.0.667 (Edgar 2010). Primers were removed and sequences of unexpected length were discarded using Cutadapt v2.3 (Martin 2011) so that only reads with a maximum deviation of 15 bp were used for further analyses (Minmax). To remove the reads with an expected error of > 0.5, the module U_max_ee was used. To normalize data, subsampling to the lowest read number per sample after merging replicates was performed to 33,523 reads using U_subset. Singletons were removed before clustering the sequences using Uparse (U_cluster_otus) with ≥ 97 % similarity into OTUs. The dereplicated sequences, including singletons, were then mapped, with a similarity of ≥ 97 %, to the generated OTU dataset to maximize the number of reads retained. OTUs with a minimal read abundance of 0.01 % in at least one sample were retained for further analyses while other OTUs were discarded. In total 8,160,330 reads that clustered into 15,812 OTUs were retained. Sequences were compared to the BOLD database sequences using BOLDigger V1_4 (Buchner et al. in review). For the whole dataset a similarity threshold of 85 % to a reference sequence in BOLD was required to include the OTU into further analysis. For taxonomic assignment of macrozoobenthic organisms a similarity of >90 % to a reference sequence in BOLD was needed.

OTU and read table analyses were performed using R Studio v.3.5.3 and figures were created using the package ggplot2 (Wickham 2009). Venn diagrams were constructed using http://bioinformatics.psb.ugent.be/webtools/Venn/. The vast majority of reads were assigned to non-arthropod taxa for the 102 samples analyzed from site 54, especially bacteria and protists (see Supplementary Fig. 1). This information was used here as one basis to design macroinvertebrate, in particular insect-specific primers.

### Primer design

As reference data we used PrimerMiner v0.18 (Elbrecht and Leese 2017b) to download COI sequences (and full mitogenomes) for 15 important freshwater macroinvertebrate orders from BOLD and NCBI, as well as their mitogenomes if available, as listed in Elbrecht & Leese (2017b) and possible non-target groups (see Supporting data A1). In addition, non-target data metabarcoding sequences were clustered into OTUs with a 3 % threshold. The most abundant non-target OTUs identified from the 102 samples collected over 15 months (i.e., algae, bacteria, fungi with a proportion of at least 1 % of total read abundance and a similarity to a sequence from BOLD or NCBI of at least 98 %) were aligned together with OTUs from the 15 MZB orders using MAFFT v7 (Katoh and Standley 2013) in Geneious v2019.2.1. The alignment was manually inspected for conserved, diagnostic bases only present in the MZB orders but not in sequences of non-target organisms. The positions of primer bases were named relative to a reference sequence of *Drosophila yakuba*, GenBank accession number X03240 (Clary and Wolstenholme 1985).

### *In silico* evaluation

The new EPTDr2n primer set was evaluated *in silico* using PrimerMiner v0.21. As COI alignments, again the 15 most important freshwater macroinvertebrate orders were used (see above). To evaluate the amplification efficiency of non-target organisms, OTU sequences from BF2BR2_OTU_table_raw_reads.csv were aligned using MAFFT as above. The primer binding regions for fwhF2 and EPTDr2n were extracted from the alignment and *in silico* evaluated using PrimerMiner on default settings. Penalty scores were grouped on Phylum level (minimum match to reference database 90 %) using the information encoded in the sequence IDs. Non-target organisms were here defined as OTUs that do not match to Arthropoda, Annelida, or Mollusca. R scripts used are available (Supplementary data A1). OTU tables were extracted as fasta files for each order using an R script and evaluated *in silico* with default parameters.

#### In vitro primer evaluation

The newly developed primer was tested using two different test cases: i) 10 samples from the biweekly long-term data series from Kinzig site 54, and ii) 10 samples from the River Kinzig network sampling (see Table S1). DNA extraction was similar to the approach outlined above. A two-step PCR was used for the newly developed primer combination fwhF2/EPTDr2n and as a comparison we used the short primer pair fwhF2/fwhR2n (Vamos et al. 2017). For the fwhF2/EPTDr2n and fwhF2/fwhR2n combinations, four length-varying primers were used in the first step with two replicates per sample, with each of them having a universal tail attached (Figure S1). Length variation was due to inline shifts (0-3 Ns) between the universal tail and the primer sequence to maximize diversity (Elbrecht and Leese 2015). For the second step, primers matching the universal tail with an i5/i7 index and P5/P7 Illumina adapter attached were used. Annealing temperature of the newly developed primer EPTDr2n together with primer fwhR2n was estimated by running a step-down gradient PCR (first 8 × 60 °C followed by 32 × 48 – 58 °C) using bulk and eDNA samples. For the first step a step-down gradient PCR with 30 cycles (6 × 60 °C for the annealing temperature and 24 × 54 °C for fwhF2/fwhR2n and 24 × 48 °C for fwhF2/EPTDr2n) was conducted, followed by the second-step PCR with 15 cycles and an annealing temperature of 60 °C. Some samples were run with 23 cycles in the second step, due to weak bands on the gel. Purification, left-sided size selection (0.78x), sequencing and sequence analysis was conducted as described above.

#### Comparison to morphological data

Classical morphological MZB taxa lists retrieved from LTER monitoring samples of up to 22 sites over the past 20 years from the River Kinzig (RMO) as well as data from a holistic RMO biodiversity database (https://rmo.senckenberg.de/search/home.php) that additionally includes data from other monitoring activities like the Water Framework Directive (WFD) monitoring were compared to taxa lists retrieved from eDNA samples using different primers. Comparisons were done at four different levels: Comparisons with 1) long-term morphological data available for an LTER monitoring site next to study site 54 (W1), 2) five LTER monitoring sites within a radius of 5 km of site 54, 3) with all LTER 22 monitoring sites of the RMO, and 4) the entire RMO database covering all monitoring activities in the RMO (e.g., WFD monitoring) (Fig. 1). Data from the ten river Kinzig sampling sites were compared similarly. For this comparison, only taxonomic names and not OTUs were considered. I.e., OTUs not matching at least at family were discarded. Different OTUs assigned to the same species were merged. This step is similar to Elbrecht et al. (2017) in order to allow for comparison between the two approaches. Taxa with ambiguous assignments (e.g. *Sericostoma personatum/flavicorne*, *Baetis* cf. *scambus*) were reduced to genus level.

An overview of the general workflow followed in this study is given in Fig. 2.

**Figure 2:**
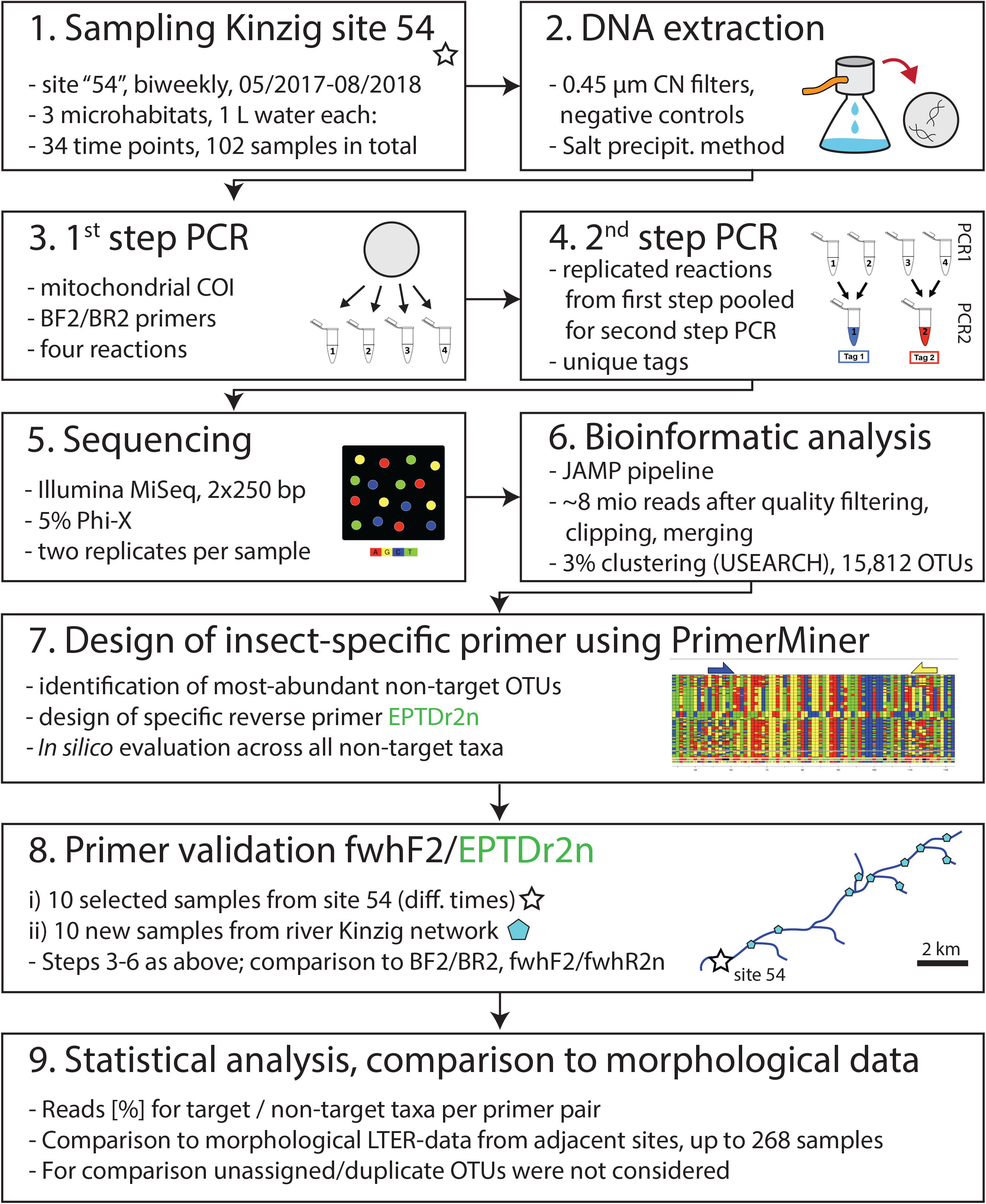
Workflow of the present study.

## Results

### Primer design and in silico evaluation

All analyzed primers are located within the classical Folmer COI-Fragment (Folmer et al. 1994)(Fig. 3A). No fully diagnostic positions were found comparing macrozoobenthic target and non-target taxa. However, two adjacent positions in the alignment (COI pos. 1979 and 1982, i.e. last and fourth last bases, Fig. 3C) differed to a large extent between target and non-target taxa. These differences were even more obvious when only considering the abundant diatom (Stephanodiscacaeae) and bacterial OTUs identified from the 102 biweekly samples analyzed with BF2/BR2 (Figure S2). This region was chosen to design a new primer EPTDr2n that can be coupled with one of the universal forward primers (e.g. fwhF2 or BF2, see Tab. 1). *In silico* analysis supports a good performance, i.e. high similarity, of fwhF2 primer across phylogenetically distinct groups with the exception of Actino- and Proteobacteria (Fig. 3B). The new reverse primer EPTDr2n had higher penalty scores for both target and non-target taxa compared to fwhF2. However, penalty scores for abundant non-target freshwater taxa (diatoms, bacteria, fungi) were substantially higher and very low for freshwater arthropods with the exception of some trichopteran genera and isopods. Trichopteran target taxa having higher penalty scores belonged to *Hydropsyche, Sericostoma, Lype*, and *Rhyacophila*, which convergently showed the same base as non-target taxa.

**Table 1.**
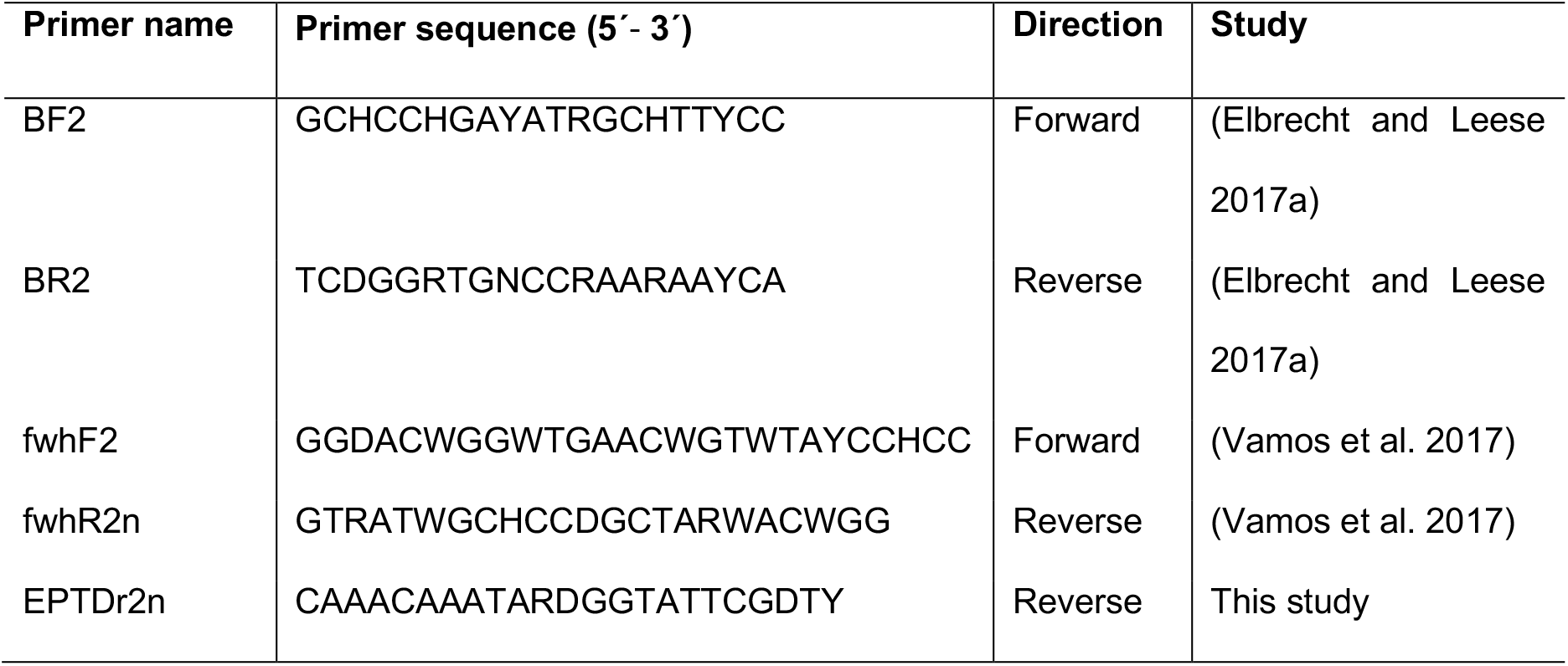
COI primers used in this study, including the newly developed EPTDr2n primer combined with forward primer fwhF2 in this study. Further primers tested that did not work are listed in Table S2.

**Figure 3:**
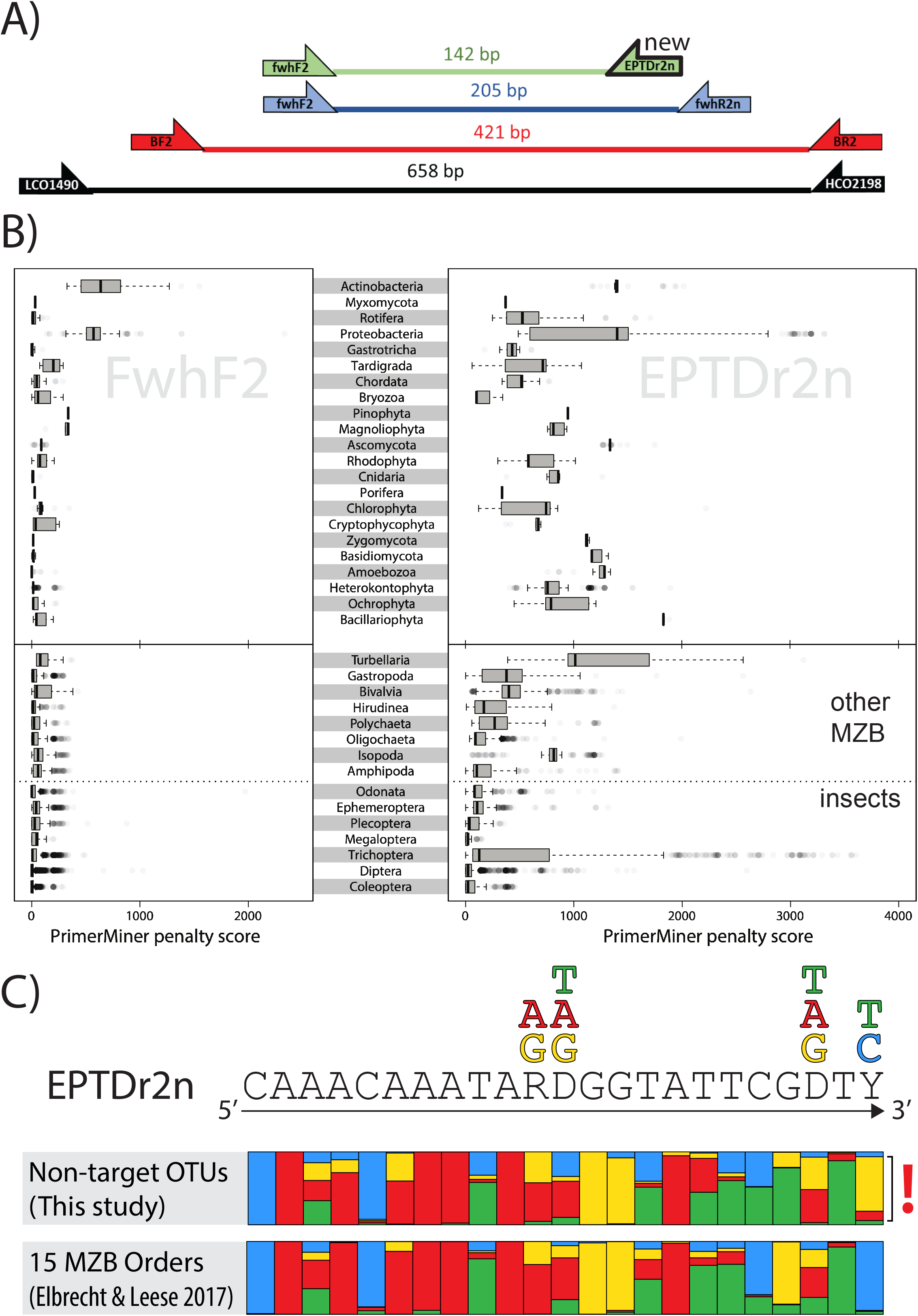
*In silico* primer evaluation. A) Overview of COI primers used including the newly developed EPTDr2n primer that was applied together with fwhF2. LCO1490 and HCO2198 (Folmer et al. 1994) primers were not used but shown for reference. B) Penalty scores for the new primer combination fwhF2 (left) and EPTDr2n (right). Results are listed for macrozoobenthic orders (MZB, bottom) and non-target groups (top). The higher the penalty score calculated using PrimerMiner (Elbrecht and Leese 2017a), the worse the primer is expected to perform. C) PrimerMiner plots of the EPTDr2n primer binding site, for 15 macrozoobenthic orders from (Elbrecht and Leese 2017a) and the non-target taxa. The quotation mark highlights the proportion of taxa omitted by choosing the the last 3’ wobble base ‘Y”.

### In vitro evaluation

#### Site 54

Sequencing was successfully conducted for BF2/BR2, fwhF2/fwhR2n and the new fwhF2/EPTDr2n primer combinations. Number of reads for primer pair BF2/BR2 ranged from 23,757 to 29,928, for fwhF2/fwhR2n from 30,683 to 32,091 and for fwhF2/EPTDr2n from 33,091 to 33,376 per sample after excluding OTUs with read numbers <0.01 % per sample. For BF2/BR2, the vast majority of reads was assigned to diatoms (56.66 %), followed by bacteria (21.98 %), fungi (5.79 %), and other algae (5.54 %). Arthropods had a read abundance of 5.38 % (see Fig. 4). Total OTU diversity was highest for BF2/BR2 (3058), moderate for fwhF2/fwhR2n (2063), and lowest for the more specific primer combination fwhF2/EPTDr2n (845) (see Supplementary data A1).

**Figure 4:**
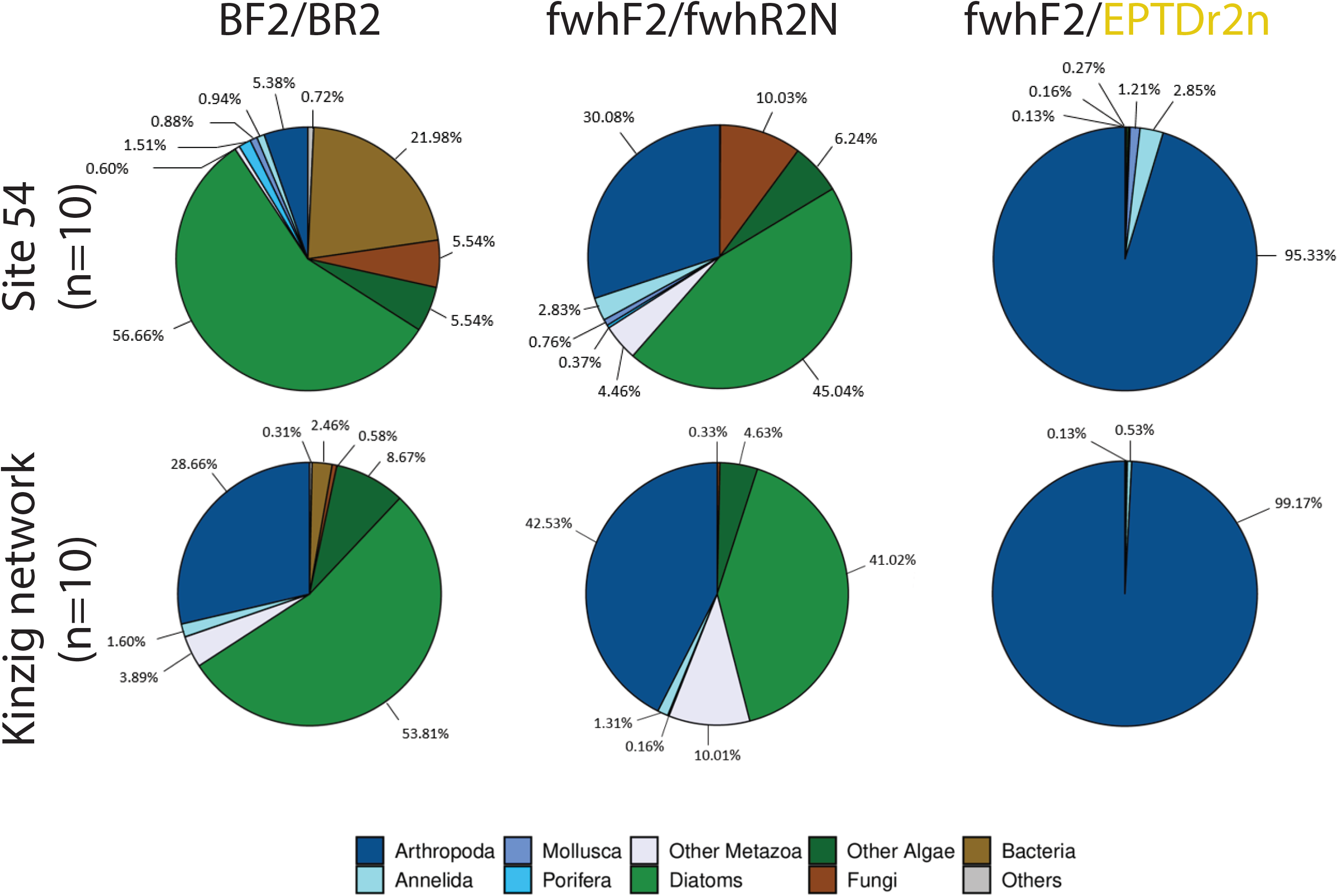
*In vitro* primer evaluation using ten 1-L samples from the biweekly RMO monitoring site 54 (top) and for ten 1-L samples from river Kinzig network (bottom). Pie charts show proportion of reads assigned to phylogenetically distinct groups (see legend) using primer combinations BR2/BR2 (A), fwhF2/FwhR2n (B), and fwhF2/EPTDr2n (C).

#### River Kinzig network

Final number of reads for primer pair BF2/BR2 ranged from 28,780 to 31,019, for fwhF2/fwhR2n from 31,849 to 33,194, and for fwhF2/EPTDr2n from 33,135 to 33,437 per sample after 0.01 threshold filtering. For BF2/BR2, the vast majority of reads were assigned to diatoms (53.81 %), followed by arthropods (28.66 %), and ‘other algae’ (8.67 %) (Fig. 4). OTU diversity was highest for BF2/BR2 (2150), moderate for fwhF2/fwhR2n (1197), and lowest for the more specific primer combination fwhF2/EPTDr2n (599) (see Supplementary data A1).

Across all 20 samples from site 54 and the river Kinzig network the number of eDNA reads targeting macrozoobenthic taxa was highest for the new primer combination fwhF2/EPTDr2n and lowest for BF2/BR2 (Fig. 5A). For the highly degenerate universal primers BF2/BR2 target read proportion was typically below 5 % and had a maximum of 34.07 % of the reads (site 5). For primer pair fwhF2/fwhR2n the number of reads of target taxa varied tremendously from below 5 % at several sites to 96.57 % at site 5 (Fig. 5A). Highest read numbers were typically assigned to dipterans, especially chironomids, independent of the primer set, but also a few oligochaete taxa (Figure S4, Supplementary data A1, B1). While primer combinations BF2/BR2 and fwhF2/fwhR2n had the overall highest OTU numbers, the number of macroinvertebrate OTUs was highest for the new primer combination fwhF2/EPTDr2n and lowest for BF2/BR2 (Fig. 5B). Interestingly, at site 5 the novel primer combination recovered a much larger proportion of OTUs compared to fwhF2/fwhR2n (114 vs. 21) despite both having similar number of reads assigned to macroinvertebrates. Here, for fwhF2/fwhR2n many reads were assigned to one chironomid OTU (genus *Rheotanytarsus*, OTU 7, 31,417 reads), whereas the new primer combination only recovered this taxon (as well as many more) with a moderate number of reads (OTU 220, 271 reads). A clear outlier site in terms of detected OTU number was site 16 from the Kinzig network. This site was sampled during the flooding season not in the river but the adjacent flooded riparian vegetation and had many reads assigned to on-target taxa but extremely few OTUs detected with all primer pairs. Almost all reads of the new primer combination were assigned to one chironomid OTU (OTU 8) of the genus *Procladius* (BOLD early release data). All other 10 OTUs found in this sample were mostly (semi-)terrestrial taxa with very low read abundances (see Supplementary data A1, B1).

**Figure 5:**
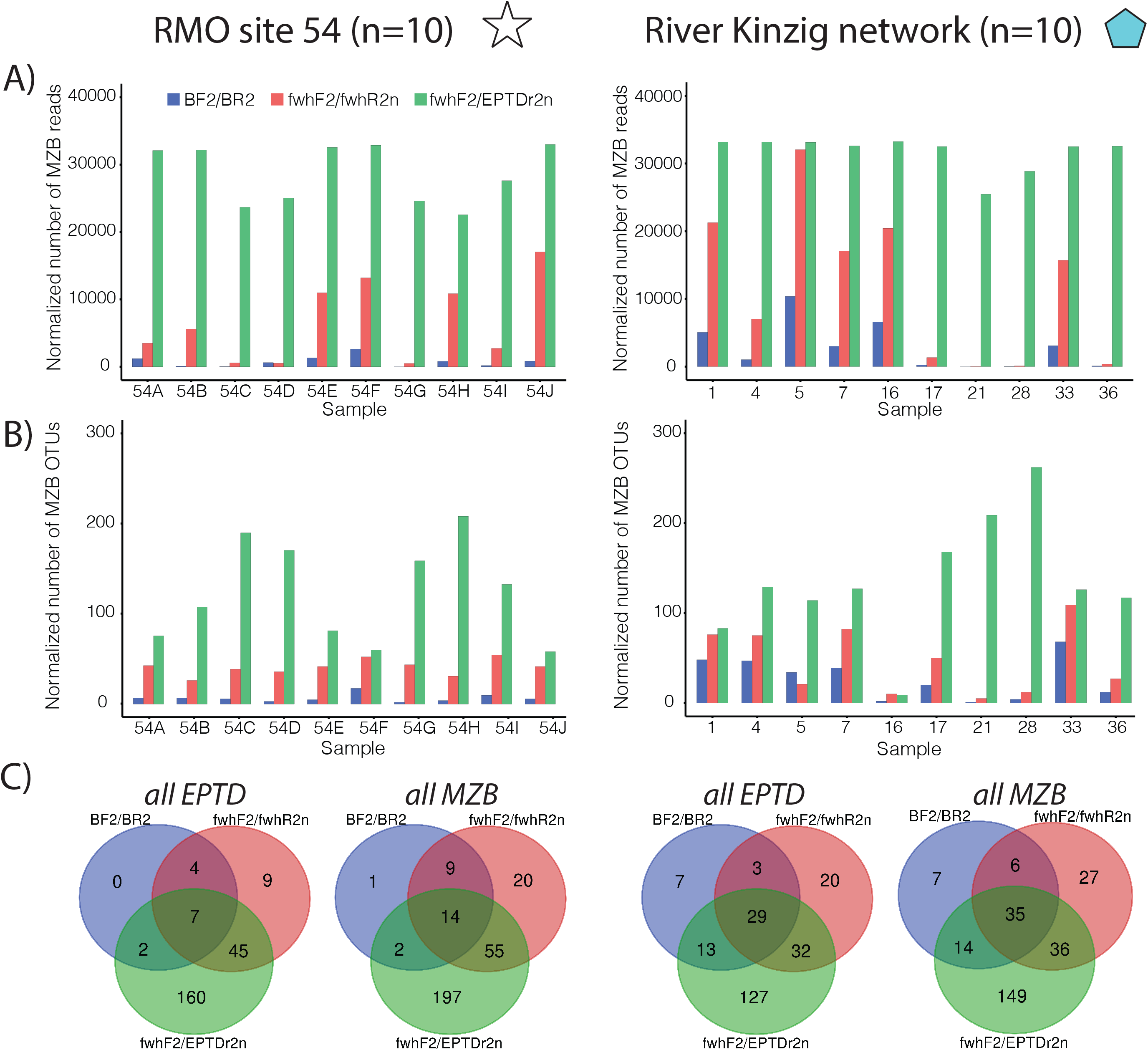
Comparison of primer performance for the two in vitro analysis test cases. A) Normalized number of reads per primer pair that target macrozoobenthic taxa (MZB). Data sets were all sub-samples to 33,523 reads to allow for comparability, B) Normalized number of OTUs per primer pair that target MZB taxa. Different samples (x-axis) represent different time points (site 54, left) and different sites (river Kinzig network, right). C) Overlap of identified OTUs for all ephemeropteran, plecopteran, trichopteran, and dipteran (EPTD) taxa as well as all MZB taxa. Note: Site 16 from river Kinzig network was an eDNA sample obtained from a floodplain.

Taxon overlap among primer combinations for ephemeropterans, plecopterans, trichopterans, and dipterans (EPTD), which are of special interest for regulatory biomonitoring and bioindication, showed that only a few EPTD taxa were exclusively found by primer combination BF2/BR2 (0 and 7 for site 54 and Kinzig network, respectively, Fig. 5C). Primer combination fwhF2/fwhR2n detected a moderate number of exclusive EPTD taxa (9 and 20, respectively) and shared the majority either with both or at least with the fwhF2/EPTDr2n primer combination. The largest proportion of exclusively detected EPTD taxa was found with the new, i.e. fwhF2/EPTDr2n primer combination (160 and 127 exclusive EPTD taxa, respectively).

When performing the comparison across all macrozoobenthic taxa that are part of the German regulatory operational taxa list, the same trend was found, yet numbers were slightly increased (Fig. 5C).

### Comparison to morphological data

For the comparison with morphological data, only OTUs with assigned taxonomy (similarity >90 %) were considered. In both *in vitro* comparisons, BF2/BR2 performed consistently worst in terms of number of identified taxa (family, genus, and species level) as well as in terms of shared taxa with morphotaxonomic lists (Tab. 2). The only four exceptions were species-level data comparison for the Kinzig network, where BF2/BR2 and fwhF2/fwhR2n performed similarly (see also Fig. 6). Primer combination fwhF2/fwhR2n performed better in 20 out of 24 and the new primer combination fwhF2/EPTDr2n best in all cases terms of total number of taxa. Number of species detected with the new primer from only 20 samples was already higher than the total number reported by expert morphological data at all 22 RMO long-term sampling site samples collected for up to two decades (n = 263, Fig. 6). In four of the 24 comparisons (Tab. 2) the number of taxa shared between eDNA and morphological datasets was slightly higher for fwhF2/fwhR2n. Here, especially trichopteran species diversity was lower for fwhF2/EPTDr2n.

**Table 2:** Comparison of long-term taxa lists generated through morphotaxonomic assessment and 10 eDNA samples obtained from site 54 (left) and the ten eDNA samples from the river Kinzig network (right). Rows list different sampling sites / site combinations of morphologically identified specimens. W1 is next to the site analyzed by eDNA (biweekly site 54, Fig. 1). Sites W1, W2, S1, S2 and O1 represent all samples (n=66) in a 5 km radius around site 54. RMO MZB stands for all samples (n=263, from 22 sites) surveyed in the course of LTER monitoring for up to 20 years. The BioDiv database covers all benthic invertebrate records (n=926) within the RMO (including e.g. data from the Water Framework Directive monitoring of local studies but also terrestrial assessments). The table is split according to three taxonomic assessment levels: species (top), genus (middle), family (bottom). The numbers in the cells, separated by ‘/’ indicate i) taxa found by both methods / ii) only DNA metabarcoding / iii) only morphology for the three primer pairs (columns).

| <u>**Species level**</u> | **Site 54 (biweekly), n = 10** |  |  | **River Kinzig Network, n = 10** |  |  |
| --- | --- | --- | --- | --- | --- | --- |
| **Morphological data** | **BF2 / BR2** | **FWH2** | **EPTDr2n** | **BF2 / BR2** | **FWH2** | **EPTDr2n** |
| **W1** | 13/24/60 | 27/110/46 | 24/254/49 | 15/133/58 | 22/118/51 | 19/239/54 |
| **W1, W2, S1, S2, O1** | 14/23/97 | 32/105/79 | 28/250/83 | 20/128/91 | 27/113/84 | 23/235/88 |
| **RMO MZB all sites** | 11/26/142 | 35/102/118 | 40/238/113 | 29/119/124 | 36/104/117 | 42/216/111 |
| **BioDiv database** | 13/24/590 | 55/82/548 | 73/205/530 | 39/109/564 | 47/93/556 | 67/191/536 |
| **<u>Genus level</u>** |  |  |  |  |  |  |
| **W1** | 12/21/49 | 28/84/33 | 28/153/33 | 21/82/40 | 27/78/34 | 28/134/33 |
| **W1, W2, S1, S2, O1** | 13/20/78 | 35/77/56 | 36/145/55 | 27/76/64 | 33/72/58 | 34/128/57 |
| **RMO MZB all sites** | 14/19/115 | 40/72/89 | 45/136/84 | 33/70/96 | 44/61/85 | 50/112/79 |
| **BioDiv database** | 13/20/285 | 47/65/251 | 56/125/242 | 37/66/261 | 43/62/255 | 53/109/245 |
| **<u>Family level</u>** |  |  |  |  |  |  |
| **W1** | 17/2/35 | 30/24/22 | 31/25/21 | 22/15/30 | 31/15/21 | 31/24/21 |
| **W1, W2, S1, S2, O1** | 17/2/51 | 34/20/34 | 36/20/32 | 23/14/45 | 34/12/34 | 35/20/33 |
| **RMO MZB all sites** | 17/2/69 | 37/17/49 | 44/12/42 | 26/11/60 | 39/7/47 | 43/12/43 |
| **BioDiv database** | 15/4/92 | 36/18/71 | 45/11/62 | 25/12/82 | 36/10/71 | 43/12/64 |

**Figure 6:**
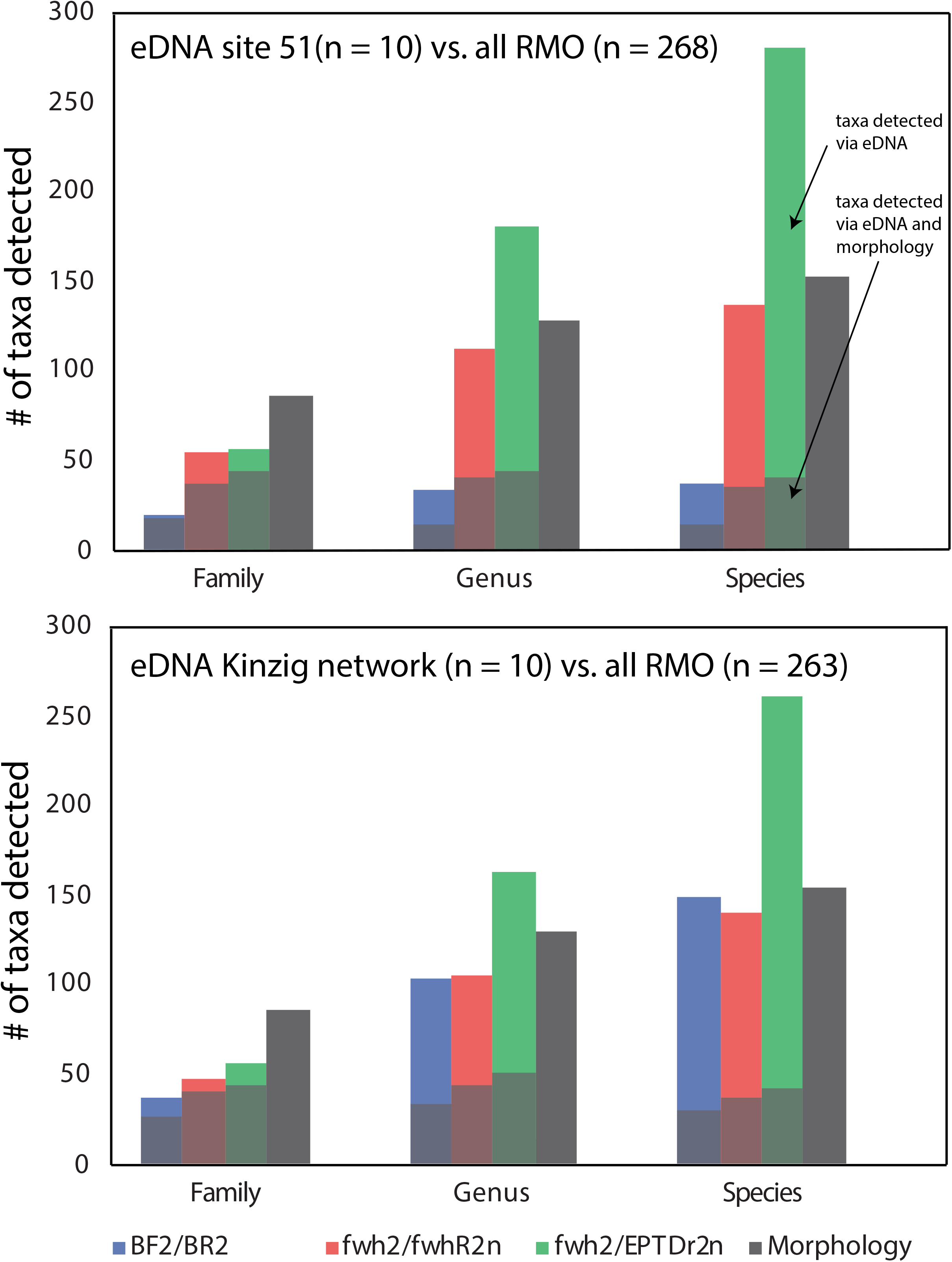
Comparison of detected taxa between eDNA and morphological assessments. eDNA based taxa lists for three primer combinations are for the 10 samples collected at site 51 (top) and the 10 river Kinzig network samples to 268 samples collected over two decades in the Kinzig catchment (grey bar). The shaded portions in the columns indicate the number of taxa shared between eDNA and morphology-based assessment.

#### Site 54

The ten samples collected at site 54 were compared the adjacent long-term monitoring site W1 (taxa list compiled over two decades of sampling, n = 28 individual samplings, reported 73 species, 61 genera, and 52 families). Across all primer pairs, the number of shared species between the two methods was below 40 %, and highest for primer pair fwhF2/fwhR2n (27 out of 73 species shared). However, with the exception of BF2/BR2, a higher number of macroinvertebrate species was reported for primers fwhF2/fwhR2n (137) and fwh2/EPTDr2n (278). This included especially many insect species (Supplementary data B1). Number of taxa exclusively reported via morphology was larger than the fraction of taxa identified with both approaches. Using the new primer combination, dipterans, but also coleopterans, plecopterans, ephemeropterans, and oligochaetes had greater taxa numbers compared to classical morphotaxonomic long-term data (Supplementary data B1, Figure S4). The total number of shared taxa between eDNA and morphotaxonomic taxa lists increased when considering taxa lists obtained from sites in a 5 km radius around site 54 (n = 66 samplings), all 22 Rhine-Main-Observatory long-term monitoring sites (n=263 samplings), or the full Biodiversity database (n = 926 assessments that includes also lentic and few terrestrial taxa, see Table 2).

#### River Kinzig network

Patterns of taxon diversity among primers was similar to the patterns reported for the 10 site 54 samples. However, shared species between eDNA and morphology were mostly lower given that the sites were further upstream (Fig. 1). Site 16, which was in the flooding zone, was a clear outlier and had very low macrozoobenthic OTU numbers (Table S1, Supplementary data B1).

## Discussion

### *In silico* analysis

We used the idea of region and ecosystem-specific primers to maximize DNA metabarcoding efficiency proposed by Elbrecht and Leese (2017a) to identify and then omit non-target taxa amplification using eDNA. eDNA metabarcoding data available from the 102 samples obtained at an LTER site (site 54) from river Kinzig allowed us to identify the most abundant non-target taxa throughout a 15-months period using BF2/BR2.

Especially Stephanodicaceae (diatoms) dominated read abundances. *In silico* analysis revealed that they differed from most ingroup taxa at position 1979 of the alignment by having either a “C”, whereas ingroup taxa had primarily a “G” (Figure S3). Few exceptions were in particular flatform, crustacean and bivalve taxa that also sometimes had an “A”. Turbellaria convergently also had sometimes a “C” like the diatoms. Our decision for a 3’ pyrimidine base, “Y”, restricted the amplification of the respective flatworm taxa but promised to avoid the abundant diatoms. The second most important position to promote amplification of ingroup taxa was at position 1982, here the trade-off was more difficult because an “A” was present in all non-target taxa but in very few of the target taxa. However, isopods as well as few trichopteran taxa (*Rhyacophila, Hydropsyche, Lype*, and *Sericostoma*) were identified to convergently share this “A”. As these taxa had a match at the last 3’-base, we accepted this mismatch to avoid higher non-target taxon binding affinity. Also, some gastropods and hirudinean taxa showed different bases at the second and third-last position,

The global *in silico* analysis performed without special emphasis on target OTUs detected at site 54 supported the general specificity of the primer, suggesting that it might not only be applicable to non-target taxa only occurring in that region. However, the analysis also again highlighted the difficulty of finding suitable primer regions that clearly separate distinct target from non-target groups (Clarke et al. 2014, Sharma and Kobayashi 2014). The global *in silico* analysis also confirmed that some taxa of Turbellaria, Mollusca, Trichoptera, and Isopoda had higher penalty scores suggesting that they might be underrepresented due to primer bias. The newly designed primer EPTDr2n was located more in the center of the ‘Folmer fragment’ compared to fwhR2n (Fig. 3). The decision to use it together with fwhF2 and not BF2 was because fwhF2 showed a particularly good match for all target groups and because the amplified region was shorter (142 bp). eDNA shed from target taxa can be degraded quickly. While the power to discriminate species decreases with shorter COI length it is still >90 % of the cases (Meusnier et al. 2008).

### *In vitro* analysis

Analysis with universal primer pair BF2/BR2 recovered mostly non-target taxa, especially diatoms, leaving only a ‘watered-down’ number of reads and OTUs of target taxa in eDNA samples, as reported for other studies using universal COI primers (Deiner et al. 2016, Macher et al. 2018, Beentjes et al. 2019, Hajibabaei et al. 2019a). This target eDNA dilution problem was consistently observed across all 20 samples analyzed as part of the two *in vitro* studies, i.e. the ten samples from site 54 (2017/2018) and the ten samples from the Kinzig network (2019). Dilution of target eDNA was less pronounced for the shorter universal freshwater macroinvertebrate primer-pair fwhF2/fwhR2n, where on average 37 % of the reads were assigned to macrozoobenthic target taxa. However, the new primer combination fwhF2/EPTDr2n completely changes this picture by yielding on average >99 % of the reads on macrozoobenthic taxa and, importantly, much greater target OTU numbers. The latter becomes obvious when comparing eDNA results from site 54 to long-term data generated for site W1, for which 117 taxa were identified by classical morphotaxonomy, whereas 351 were detected with eDNA metabarcoding. When restricting the analysis to the years 2017/2018, the difference became even larger (Figure S5, Supplementary data B2). Especially for trichopterans, plecopterans, and dipterans the number of taxa identified morphologically decreased considerably and is even lower than the number of taxa identified with the universal fwhF2/fwhR2n primer pair. In addition, the majority of taxa recovered with eDNA metabarcoding was identified to species level and only few to family level in comparison to taxa found at site W1. This is not unexpected given that a higher taxonomic level for macroinvertebrates is often kept to avoid species misidentification especially for small specimens (Haase et al. 2010). Even when considering the taxon number for all 263 samples obtained from 22 long-term monitoring sites in the past 5 - 20 years, the number of macrozoobenthic species and genera detected is greater for the eDNA samples of the two case studies compared to the morphologically retrieved. It must be noted, however, that family diversity was higher using the long-term morphological data. This is owed to the fact that i) primer bias omits taxa such as some Turbellaria and Mollusca, and ii) a much smaller temporal and spatial sampling is captured with the few eDNA samples. Overall, the dominating taxa in eDNA metabarcoding were insects, especially dipterans (chironomids, simuliids), ephemeropterans, coleopterans, and plecopterans but also oligochaetes (see Figure S4). The high proportion of reads and OTUs assigned to chironomids (Supplementary data B1) are expected as non-biting midges are known to dominate freshwater ecosystems in terms of abundance and diversity (Pinder 1986, Armitage et al. 1995). Applying bulk metabarcoding Beermann et al. (2018) found 183 chironomid OTUs in a small German low-mountain stream using the same 3 % distance threshold as applied here, which is similar to the high OTU diversity found here. While the new primer combination comes with a substantial gain in arthropod, in particular insect diversity, the increased specificity also has a down-side because using the new EPTDr2n also excluded some derived arthropod target taxa that convergently showed the same base as the non-target taxa, e.g. trichoptera genera *Rhyacophila, Hydropsyche*, and the isopod *Asellus aquaticus*. These were captured with primer pairs fwhF2/fwhR2n (and in part with BF2/BR2). Likewise, flatworms, and molluscs were also not, or not reliably detected with the new primer combination. For molluscs and crustaceans, specific primers targeting the 16S gene have been proposed (Klymus et al. 2017, Komai et al. 2019) and should be considered as a complement depending of the goal of a study.

### Considerations for macroinvertebrate bioassessment

The choice of primers is always a difficult quest and depends on the aim. However, no ‘one-fits-all’ solution exists (Clarke et al. 2017, Elbrecht and Leese 2017a, Taberlet et al. 2018, Grey et al. 2018, Hajibabaei et al. 2019b, Elbrecht et al. 2019). For arthropod bulk samples, good primers exist (Elbrecht et al. 2019). If the aim of a study is to capture the greatest number of macrozoobenthic taxa, in particular arthropods from eDNA for bioassessment with one primer pair, the newly designed fwhF2/EPTDr2n combination fits that purpose. It clearly outperforms even catchment-wide macrozoobenthic species numbers (Fig. 6). However, the overlap between eDNA and morphological assessments is moderate. This may also be due to the very limited number of samples and time-points collected via eDNA. If the aim is to maximize the phylogenetic diversity captured in a sample either a more conservative marker such as 18S can be appropriate (Deagle et al. 2014, Li et al. 2018, Bagley et al. 2019), or - for metazoan taxa - it might be important to consider the use of multiple COI primers (Corse et al. 2019, Hajibabaei et al. 2019b) as with the more specific primer combination clearly a few macrozoobenthic taxa within Trichoptera, Mollusca, and Isopoda will be missed due to primer bias. It should be noted that also for broad eukaryotic markers the diversity of macrozoobenthic species can still be diluted and read numbers dominated by meiofaunal groups that life in the habitat (Rotifera, Copepoda, see Li et al. 2018) but which are usually not part of the regulatory biomonitoring.

The new primer combination developed here has successfully been tested *in silico* and used *in vitro* for samples from one German LTER site. Non-target taxa show much lower binding affinities at the 3’-end compared to most arthropod taxa in the global analysis (Fig. 3B) and thus we are confident that the same primer will work also in other regions / aquatic habitats and also on sample preservative liquid and completely homogenized environmental samples (Zizka et al. 2018, Beerman et al. unpublished data, Pereira-da-Conceicoa et al. 2019, Blackman et al. 2019). It can possibly also be further improved by adding further variability, yet too many degenerate bases also limit amplification success (see Supplementary Table 2 for primers not working). The increased resolution for ecologically important taxa such as chironomids and oligochaetes can be of immense relevance for biomonitoring (Milošević et al. 2013, Vivien et al. 2015, 2016, 2020, Macher et al. 2016, Beermann et al. 2018). They dominated eDNA signals here and were often site-specific. The same has been shown for meiofaunal stream biota recovered from the large river Yangtze in China (Li et al. 2018). Therefore, the drastically increased resolution of such ecologically important indicator taxa is a clear benefit when it comes to biomonitoring (Pawlowski et al. 2018). Of course, signals derived from eDNA collected from water will always integrate across a greater section of the river and thus intercalibration with traditional site-based methods is difficult. However, recent studies showed that eDNA signals can pick up very local signals despite the current regime (Macher et al. 2018, Li et al. 2018, Jeunen et al. 2019). Thus, it is worth considering it as a complementary tool for bioassessment and monitoring also of stream invertebrates.

In conclusion, our study shows that the detection of macrozoobenthic taxa from eDNA isolated from stream water is greatly increased with a new specific primer combination that avoids non-target taxa amplification. The primer and approach proposed here offer a solution to the common problem of ‘watered-down’ macrozoobenthic biodiversity in COI eDNA metabarcoding and may thus improve future biodiversity assessment and monitoring of freshwaters.

## Acknowledgments

We would like to thank Marlen Mährlein, Nathalie Kaffenberger, Cristina Hartmann-Fatu and Arne Beermann for help with sample collection or processing. Furthermore, we thank Jan-Niklas Macher for helpful discussions, Beatrice Kulawig for compiling the taxonomic data from the Rhine-Main-Observatory, and Martina Weiss for valuable comments on the structure of the manuscript. This work has been funded by a grant of the Bode Foundation to FL. This work has been conducted as part of COST (European Cooperation in Science and Technology) Action DNAqua-Net (CA15219).

## Supporting information

**Supporting information file**: pdf with supplementary tables S1 and S2 as well as supplementary figures S1 – S5.

**Supporting data A1:** R scripts for *in silico* analysis.

**Supporting data A2:** OTU and read tables.

**Supporting data B1:** Filtered taxon lists (no duplicate names, only hits with >90 % similarity to BOLD) for all site 54 and River Kinzig data sets. Also taxon lists based on morphological surveys listed.

**Supporting data B2:** Morphotaxonomic lists for site W1 for the years 2017/2018.

**Data accessibility:** All read data can be found in the Short-Read Archive, Project number: XX will be provided prior to publication XX

