## Supporting Tables and Figures for "Improved freshwater macroinvertebrate detection from eDNA through minimized non-target amplification"

**Supplementary file to the article**

### A new specific COI primer significantly improves biodiversity assessments of stream arthropod communities via environmental DNA

**Table S1.** Samples used for testing the performance of the newly designed primer EPTDr2n. Ten samples from the river Kinzig network and ten from the long-term monitoring (site 54) were used for comparison. Green for high Bacillariophyta reads, blue for high EPTD reads per sample. All information are added as supporting tables with taxonomy assigned / read numbers.

| Sample / site | Number of EPTD reads detected with BF2/BR2 | Number of Bacillariophyta reads detected with BF2/BR2 | Project | Info | Original sample name | Longitude | Latitude |
|---|---|---|---|---|---|---|---|
| 1 | 45269 | 30075 | Kinzig network analysis | Kinzig before Elmbach inflow | Ki_EBVO | 50.345883 | 9.531387 |
| 4 | 9255 | 140414 | Kinzig network analysis | Elmbach before Schwarzbach inflow | EB_SBVO | 50.361343 | 9.558884 |
| 5 | 77156 | 70135 | Kinzig network analysis | Elmbach after Schwarzbach inflow | EB_SBNA | 50.358253 | 9.550212 |
| 7 | 22474 | 53548 | Kinzig network analysis | Kinzig before Ahlersbach inflow | Ki_ABVO | 50.326005 | 9.4902 |
| 16 | 61423 | 46326 | Kinzig network analysis | Sample from high water in March | 6.2 | 50.1959672 | 9.1755142 |
| 17 | 2146 | 168829 | Kinzig network analysis | Bracht before flowing into the Kinzig | KiBrZu | 50.25465 | 9.304 |
| 21 | 114 | 125015 | Kinzig network analysis | Kinzig before Gründau inflow | KiGrVO | 50.16129 | 9.02269 |
| 28 | 295 | 136883 | Kinzig network analysis | Kinzig before Birkigsbach inflow | KiBiVOS | 50.18304 | 9.09173 |
| 33 | 24112 | 54602 | Kinzig network analysis | Grennelbach before flowing into the Kinzig | KI_GBZU | 50.34198 | 9.58398 |
| 36 | 914 | 131307 | Kinzig network analysis | Kinzig before Salz inflow | KiSaVO | 50.27585 | 9.36073 |
| 54A | 3324 | 4583 | RMO-biweekly | 19.07.2017, riverbank | RS3 | 50.132970 | 8.954320 |
| 54B | 221 | 45708 | RMO-biweekly | 02.08.2017, riverbed | MS21 | 50.132970 | 8.954320 |
| 54C | 67 | 14900 | RMO-biweekly | 25.10.2017, surface | AB3 | 50.132970 | 8.954320 |
| 54D | 1753 | 12889 | RMO-biweekly | 25.10.2017, riverbed | CH2 | 50.132970 | 8.954320 |
| 54E | 3058 | 5248 | RMO-biweekly | 08.11.2017, riverbed | AB2 | 50.132970 | 8.954320 |
| 54F | 5426 | 24392 | RMO-biweekly | 14.03.2018, riverbed | db2 | 50.132970 | 8.954320 |
| 54G | 8 | 84399 | RMO-biweekly | 28.03.2018, surface | BP7 | 50.132970 | 8.954320 |
| 54H | 43 | 23591 | RMO-biweekly | 09.05.2018, riverbed | MS2 | 50.132970 | 8.954320 |
| 54I | 133 | 8279 | RMO-biweekly | 06.06.2018, riverbed | MS23 | 50.132970 | 8.954320 |
| 54J | 3641 | 2456 | RMO-biweekly | 15.08.2018, riverbed | MS26 | 50.132970 | 8.954320 |

**Table S2:** Overview of primers with higher codon degeneracy tested unsuccessfully in this study in all pairs as well as with the fwhF2/fwhR2n primers as second primer.

| | | | | |
|---|---|---|---|---|
| EPTDf1 | GHWBHHTWRTWGAWANHGGDRC | 22 bp | Forward | This study |
| EPTDf1n | GWAGHHTWRTWGAWADHGGDRC | 22 bp | Forward | This study |
| EPTDr1 | BNDVNWYKYATRTTWATWRYDGTDG | 25 bp | Reverse | This study |
| EPTDr2 | CANACRAAWARDGGNATWHKDTY | 23 bp | Reverse | This study |

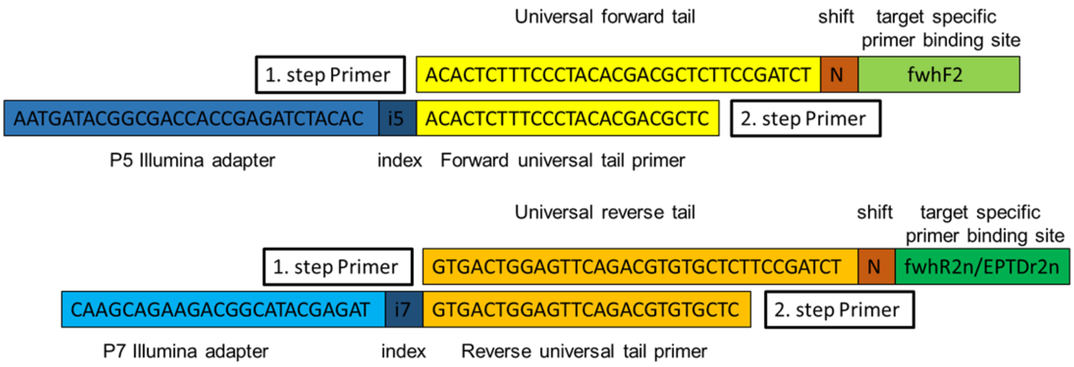


***Figure S1.*** Primer structure for the fwhF2 and EPTDr2n primers. For the first step four variants of the primer with a universal tail and an inline shift of 0-3 Ns are equally mixed. In the second step, universal primers bind to the tail of the first step primers and attach the index and Illumina adapter, needed for sequencing. 5‘ is on the left side (for all four primers shown).

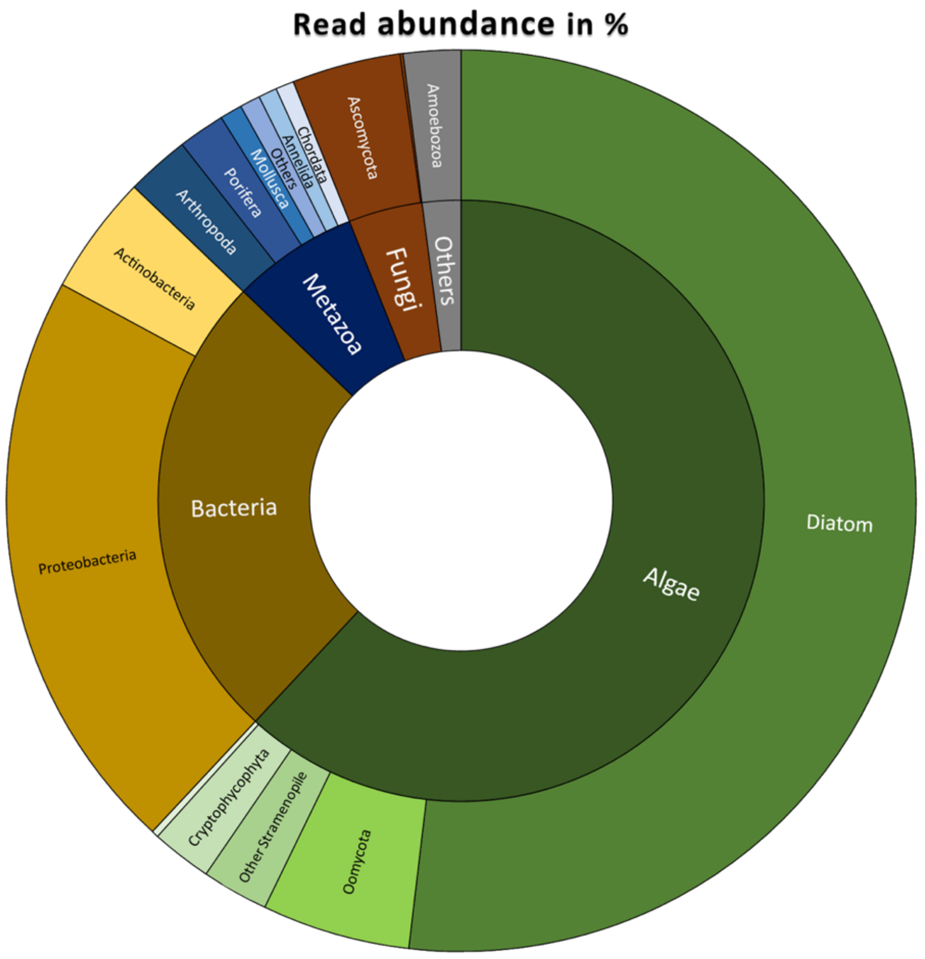


**Figure S2:** Proportion of reads assigned to different taxa using the 102 samples obtained over the 15-months sampling campaign with primers BF2/BR2.

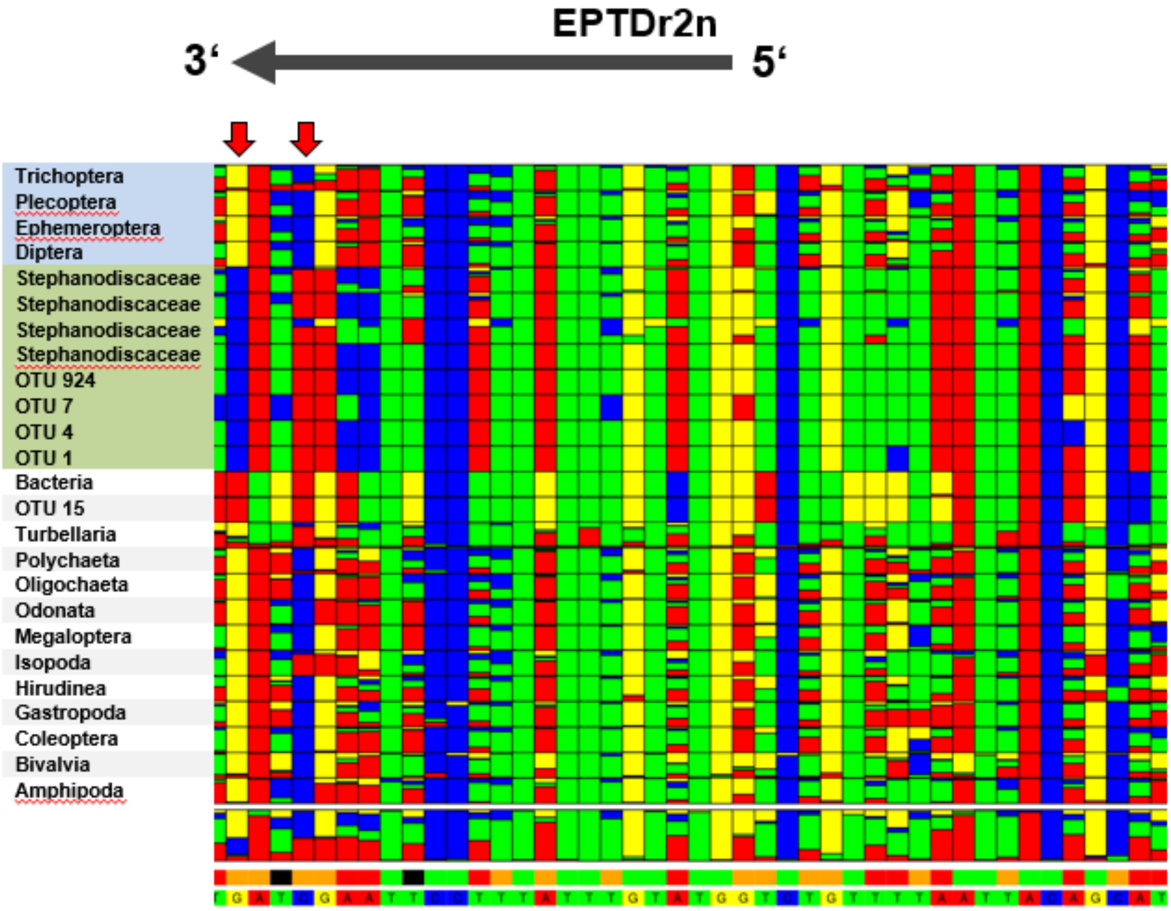


**Figure S3:** Alignment of the 15 MZB taxa (Elbrecht and Leese, 2017), Stephanodiscaceae, Bacteria sequences (extracted from BOLD and NCBI) and the most abundant non-target OTUs (1,4,7,15,924) from the 102 biweekly BF2/BR2 dataset (see Materials and Methods). Blue marks the EPTD and green the algae sequences. Alignment showing the EPTDr2n primer binding site. Positions **1979** (left) and **1982** (more to the right) in the COI fragment are marked with red arrows relative to reference sequence of *D. yakuba*, Clary & Woltsenholme (1985).

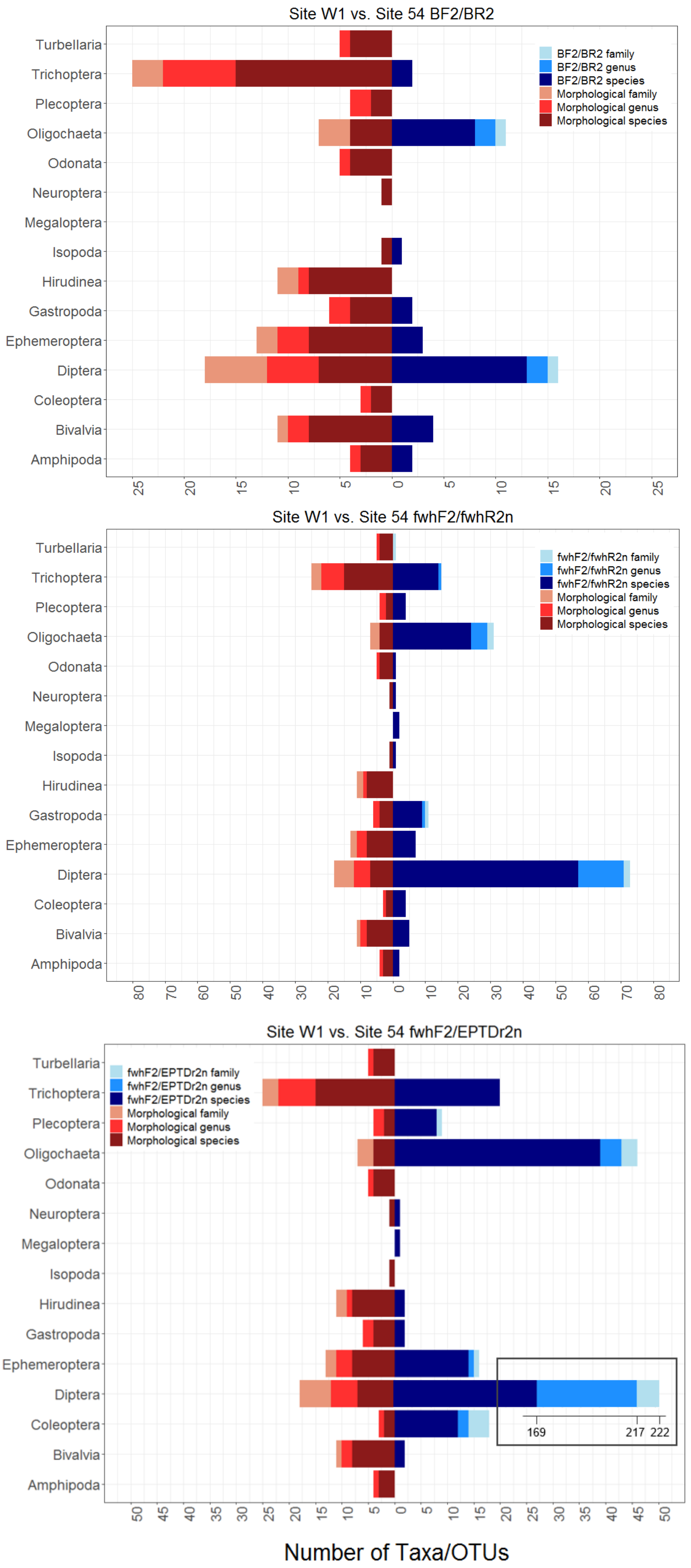


**Figure S4:** Comparison of taxa recovered via morphological methods at site W1 (n=28) and via the 10 eDNA samples from site 54 from the years 2017 and 2018 and three different primer combinations. From top to bottom: BF2/BR2; fwhF2/fwhR2n; fwhF2/EPTDr2n. OTUs assigned to species in dark blue/red, taxa assigned only to genus in middle blue/red and those only assigned to family in bright blue/red (additive).

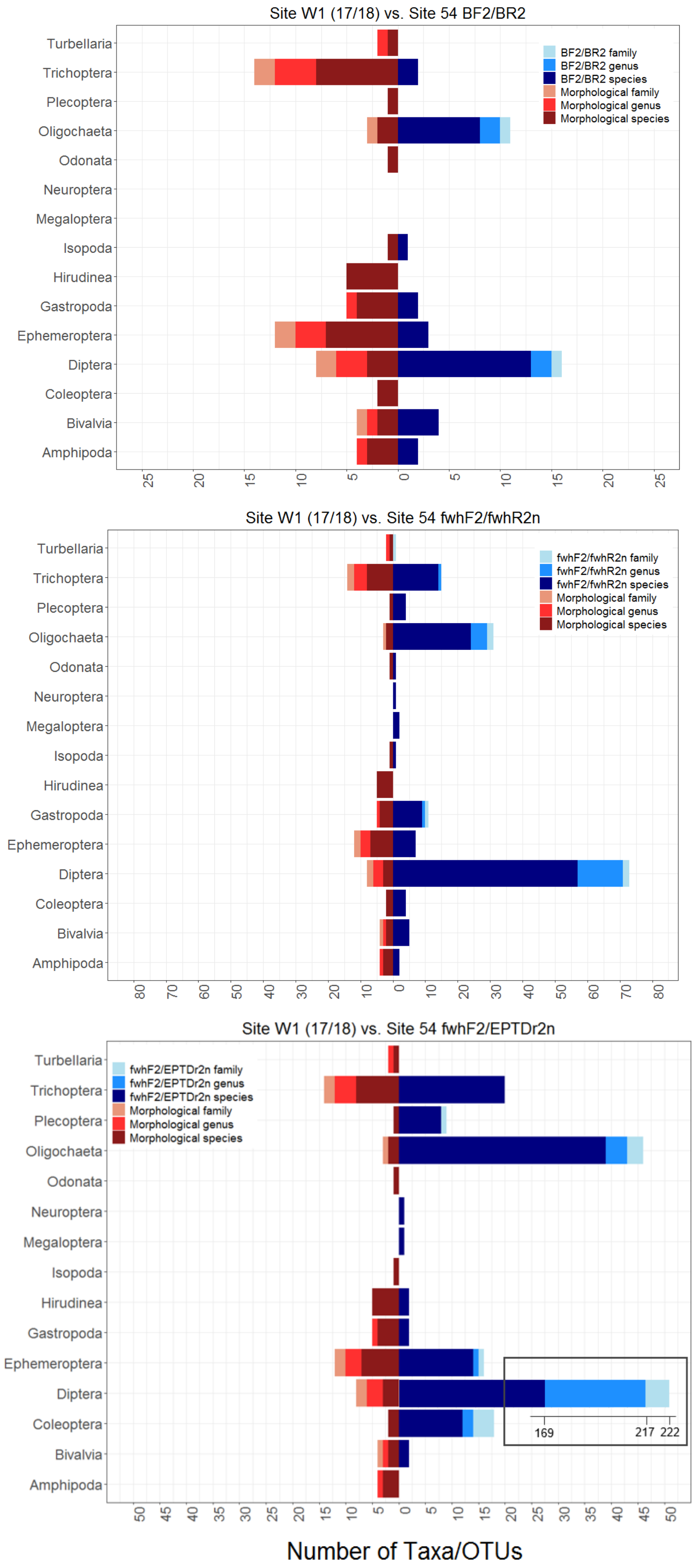


**Figure S5:** Comparison of taxa recovered via morphological methods at site W1 only from the years 2017 and 2018 and using the 10 eDNA samples from site 54 from the years 2017 and 2018 and three different primer combinations. From top to bottom: BF2/BR2; fwhF2/fwhR2n; fwhF2/EPTDr2n. OTUs assigned to species in dark blue/red, taxa assigned only to genus in middle blue/red and those only assigned to family in bright blue/red (additive).
